# The Effect of E-Cigarettes on the Human Heart Studied using Cardiomyocytes Generated from Induced Pluripotent Stem Cells

**DOI:** 10.1101/2023.04.06.535946

**Authors:** Ava T. Bhowmik, Shane R. Zhao, Joseph C. Wu

## Abstract

Although many dangerous effects of smoking e-cigarettes on lungs have come to light, limited and only qualitative efforts have been made to analyze the impact of e-cigarettes on the human heart directly. In this study, we experimentally determined e-liquid cardiotoxicity in both healthy cells and cells with long QT syndrome by treating healthy and diseased human cardiomyocytes with e-liquids with varying nicotine concentrations. These cardiomyocytes were generated from human induced pluripotent stem cells. The cardiomyocytes were divided into 5 groups, a control group and 4 test groups, each treated with e-liquid containing varying amounts of nicotine between 0% and 70%. The cells’ biological indicators such as heart rate, pulse pressure, essential protein concentration, and metabolic activity, were measured and characterized using three different functional assays: contractility, Western blot, and viability, and tracked over 4 days. The results demonstrated that acute exposure to e-liquid led to tachycardia, hypertension, decreased protein levels, and cell death. The rate of cardiotoxicity increases with higher nicotine concentrations. The basal fluid also showed non-negligible toxicity. Under identical conditions, the functionality of the diseased heart cells declined at a faster rate compared to healthy cells. Overall, this work establishes the harmful physiological effects of e-cigarettes on the human heart quantitatively.

## I. INTRODUCTION

Electronic cigarettes (e-cigarettes) have become increasingly popular with adolescents in recent years as a result of aggressive marketing schemes, false safety claims, and appealing flavors targeted towards teens from e-cigarette companies. In the past 8 years alone, e-cigarette use amongst youth has increased by 18 times (Hudson, 2022).

The anatomy of a typical e-cigarette is shown in Figure 1 (Brown, 2013). E-cigarettes contain a “juice” in a cartridge, which is referred to as e-liquid. It is composed of various carrier compounds, additives, flavorings, and nicotine. The e-liquid in the cartridge is heated in the vaporizing chamber and delivered as aerosols through the mouthpiece to the user. The battery voltage is adjustable to allow users to customize vapor discharge quantity per puff. The inhalation of this e-liquid vapor could be dangerous to pulmonary and cardiovascular health.

**Fig. 1.**
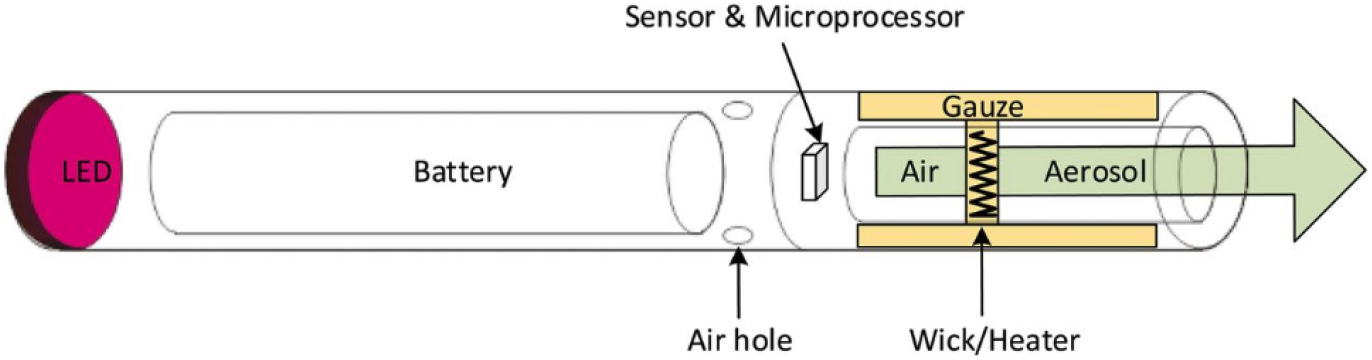
Anatomy of a typical e-cigarette (Brown, 2013). The e-liquid is vaporized in the vaporizing chamber, which consists of a nicotine cartridge, heating coil, and atomizer. This e-liquid is thought to have cardiotoxic effects on human cardiomyocytes.

Several published studies have shown that smoking e-cigarettes negatively affects pulmonary health and has carcinogenic effects on lungs (Rowell, 2015; Hecht, 2015), and impacts the cardiovascular system (Qasim, 2017). However, no direct experimental studies have been reported to quantify the physiological effects of smoking e-cigarettes on the human heart cells. As such, we decided to use cardiomyocytes generated from induced pluripotent stem cells to quantitatively determine e-liquid cardiomyopathy.

Induced pluripotent stem cells (iPSCs) are pluripotent stem cells derived from somatic cells, such as blood cells or skin cells (Ye, 2013). Since their discovery by Shinya Yamanaka in 2006 (Takahashi, 2006), numerous researchers have utilized human induced pluripotent stem cells (hiPSCs) to carry out drug discovery, health studies, and “clinical trials in a dish” (Grskovic, 2011; Karakikes, 2015; Matsa, 2016; Shi, 2017; Magdy, 2018; Fermini, 2018; Lee, 2019). In particular, hiPSCs have been used to generate cardiomyocytes to conduct cardiovascular research (Chatterjee, 2010; Matsa, 2011; Burridge, 2016; Narkar, 2017; Kitani, 2019; Wei, 2022).

In this project, the impact of e-liquid on the human heart cells was experimentally studied and quantified. In addition, we compared the magnitude of the impact between healthy and diseased heart cells. We hypothesized that even the basal e-liquid without nicotine could possess compounds that induce cardiomyopathy; higher nicotine concentrations in the e-liquid cause more rapid decline in cardiomyocyte health; and under equivalent conditions, the effect would be worse for diseased cells as compared to healthy cells.

Thus, we conducted a series of experiments using cardiomyocytes that we differentiated from two lines of hiPSCs – one healthy and one with long QT syndrome. To correlate the e-liquid’s nicotine content to cardiotoxicity, the cardiomyocytes were divided into five groups: a control group, one group treated with pure basal fluid without nicotine, and three groups each to be treated with e-liquid containing varying amounts of nicotine of 30%, 50%, and 70%, respectively. To understand the effect of e-liquid exposure on heart rate and pulse pressure, essential protein concentration, and metabolic activity, we used three different functional assays to quantify the cardiotoxicity of the e-liquids; contractility, western blot, and viability.

## II. PRIOR RESEARCH

There have been numerous human observational studies, murine experiments, and review papers on the effect of smoking cigarettes and e-cigarettes on the heart (Qasim, 2017; Papathanasiou, 2013; Lee, 2018). It has been established that nicotine has cardiotoxic effects and that smoking is a risk factor for heart attacks and myocardial infarction (Ottervanger, 1995). Cardiovascular disease is a major cause of death among smokers (Goldenberg, 2003; Qasim, 2017). Papathanasiou et al. reported that nicotine causes an acute increase in heart rate and blood pressure in adult humans (Papathanasiou, 2013).

Lee et al. reported their findings on the effect of e-cigarette smoke on a mouse heart as measured by DNA damage (Lee, 2018). In this work, to debunk the claim that e-cigarettes are noncarcinogenic, a group of mice were exposed to e-cigarette smoke. It was found that treatment with e-cigarette smoke caused damage to mouse DNA and slowed repair processes in the murine lung, heart, and bladder. However, it still remained unclear whether this cardiotoxic effect also extended to humans.

Farsalinos et al. analyzed the National Health Interview Surveys (NHIS) of 2016 and 2017 to examine whether e-cigarette use was consistently associated with coronary heart disease and myocardial infarction (Farsalinos, 2019). However, they were unable to draw a significant association between e-cigarette use and an elevated risk for coronary heart disease or myocardial infarction in humans.

Use of hiPSCs for experimental purposes was demonstrated by Wei et al. Their study established that cannabis users had a higher risk of myocardial infarction and that genistein attenuates marijuana-induced vascular inflammation (Wei, 2022).

A review paper by Cherian et al. described e-cigarette associated lung injury. It was found that vaping products gave rise to a variety of complications, including thermal injuries, life-threatening pulmonary issues, and more (Cherian, 2020). Although the pathological effects of e-cigarettes on human lungs are mostly established, there have been no systematic and quantitative studies to determine how e-liquid affects human heart cells physiologically thus far. Although much is known about smoking-induced cardiotoxicity, very little is known about vape-induced cardiotoxicity. This motivated our work to experimentally determine the effect of e-cigarettes on human cardiomyocytes derived from hiPSCs.

## III. METHODS AND MATERIALS

### A. iPSC Confluency

In this work, we passaged and cultured two lines of iPSCs until they reached 80% confluency. One line contains healthy cells and the other line contains cells exhibiting long QT syndrome.

First, for each line, we precoated two 6-well cell plates with 1.5 mL of Matrigel® per well. This forms a coating at the bottom of each well to help the newly-passaged iPSCs to attach to the plates. Then, we passaged existing cells of each line at a 1 to 12 ratio, redistributing them into 12 wells. To do this, we first digested one well of iPSCs with Gentle Cell Dissociation Reagent (GCDR). After 5 minutes, we aspirated the fluid and added 1 mL of Insulin RPMI with B27 supplement; we then fully separated the layer of cells from the bottom of the plate by pipetting up and down. Once the Matrigel®-coated plates incubated at 37°C for at least 30 minutes, we added 1.5 mL of StemMACS™ iPS brew XF to each well of the two 6-well plates. Finally, we added 80 *µ*L of the dissolved cell solution to each well. After proper settling, the plates were incubated at 37°C with 5% CO2. The iPS brew medium was replaced daily until the cells reached 80% confluency, which usually takes about 3 days to occur.

### B. Cardiomyocyte Differentiation

The process of differentiation starts after the iPSCs reach roughly 80% confluency and generally takes between 8 to 12 days. On Day 0, we replaced the iPS brew medium with 1.2 mL of 6 *µ*M CHIR99021 (CHIR). The addition of CHIR activates the Wnt signaling pathway to induce mesoderm differentiation. We diluted 0.9 mL of 10 *µ*M CHIR stock solution with 0.3 mL of -Insulin RPMI to achieve a 6 *µ*M CHIR solution. Based on the results, concentration of CHIR may require adjustment up to 8 *µ*M or 10 *µ*M to ensure proper induction of mesoderm differentiation. Mesoderm differentiation continued on Day 1. Day 2 was the recovery phase; we neutralized the CHIR with an equal volume (1.2 mL) of -Insulin RPMI. Cardiac mesoderm specification began on Day 3 and extended to Day 4. On Day 3, we replaced the medium with 2 mL of 10 *µ*M IWR stock solution diluted to 5 *µ*M using -Insulin RPMI. The IWR inhibits the Wnt signaling pathway once fully differentiated. We then neutralized the medium with 2 mL of -Insulin RPMI on Day 5 to allow for recovery into Day 6. Day 7 marks the completion of differentiation; beating cardiomyocytes may be observed at any time from days 6-11. In this experiment, we observed beating cardiomyocytes on Day 6. For the completion of the differentiation process, we replaced the -Insulin RPMI with 2 mL of +Insulin RPMI to provide nutrients to the cells. Figure 2 shows the progression of differentiation for this experiment.

**Fig. 2.**
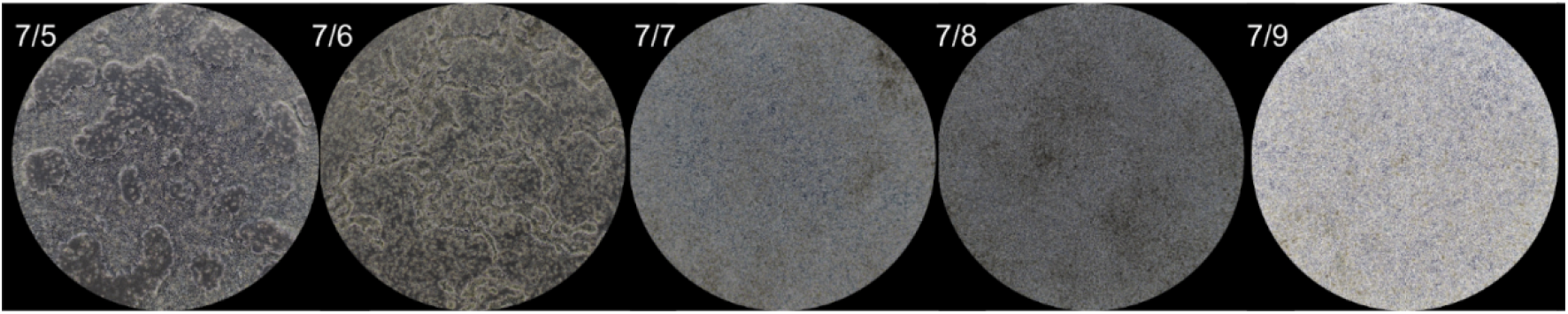
Optical microscope images depicting the progression of cell growth over 5 days after reaching full confluency. Visible morphological changes can be observed throughout the process as the cells differentiate. Beating cardiomyocytes were observed on Day 6.

### C. E-liquid Treatment

After generating the beating cardiomyocytes from iPSCs, we exposed both the healthy and diseased cells to the e-liquid. We obtained 4 different e-liquids, the basal fluid with no nicotine content, and Marcado-flavored e-liquids with 30%, 50%, and 70% nicotine concentration by volume from Red Oak.

First, we replated each cell line of the cardiomyocytes onto 96-well plates, ensuring a cell count of at least 3,000 per well and 80 wells per line. We used 16 wells as control and 16 wells each for every concentration of nicotine. To emulate the e-cigarette vaporization process and to account for potential degradation in the e-liquid components, we first boiled the e-liquid in a CFX Duet BioRad polymerase chain reaction (PCR) machine. We waited for the vapor to cool and condense before adding 10 *µ*L of condensed e-liquid at each intended concentration to the wells.

This research only tested the effects of acute (rather than chronic) exposure to e-liquid, so in subsequent weeks we replaced the medium with -Insulin RPMI every two days.

### D. Functional Assays

We utilized three different functional assays to quantify the impact of e-liquid exposure on the cardiomyocyte health: contractility, western blot analysis, and cell viability. We used the contractility assay to determine the general physiological effects of e-liquid on the heart by monitoring cardiomyocytes’ heart rate and pulse pressure after 3 days of exposure. To analyze the e-liquids’ impact at the cellular level, we tracked protein levels in the cardiomyocytes via western blot. Finally, to correlate the rate of cell death with e-liquid treatment, we observed cell viability using the Countess Cell Counting Chamber. Each data point in the results were collected and averaged over 16 wells.

1. *Contractility:* We used the Sony SI8000 Cell Motion Imaging System to determine the heart rate and pulse pressure of the cells. These markers are useful in determining cell health as well as extending cellular symptoms to real-life conditions, such as bradycardia vs. tachycardia or hypotension vs. hypertension.
2. *Western Blot Analysis:* We lysed the cells using ThermoFisher radioimmunoprecipitation assay (RIPA) buffer and centrifuged the lysates at 13,000 rotations per minute (rpm) for 15 minutes at 4 °C. The lysates were then subjected to sodium dodecyl sulfate–polyacrylamide gel electrophoresis (SDS-PAGE) using polyacrylamide gels. The western blot was done as previously described (Pillai-Kastoori, 2020; Smith, 2009). We performed immunoblotting using specific antibodies in order to evaluate the expression of different proteins.
3. *Cell Viability:* To test the cell viability, we first stained the cells with Trypan blue dye diluted at a 1:10 ratio with phosphate buffered saline (PBS) and placed the stained cells into the Countess Cell Counting Chamber slides. Then, we utilized the ThermoFisher Countess Cell Counting Chamber to characterize the percentage of cells that are viable over time.

## IV. RESULTS AND DISCUSSION

After tracking the performances of the healthy and diseased cell lines over a course of 4 days, we analyzed the results and determined the e-liquids’ cardiotoxic effects on the cardiomyocytes when they were acutely exposed. In the following sections, we discuss the experimental results from three different functional assays that we carried out: contractility, western blot, and viability.

### A. Impact on Contractility

The e-liquid’s impact on cardiomyocytes’ pulse rate, measured in beats per minute (bpm), and pulse pressure, measured in millimeters of mercury (mmHg), was studied through the contractility assay. The untreated cells served as a baseline for control; their pulse rate stayed relatively constant over the course of 4 days. However, the cells treated with e-liquid all experienced tachycardia, beating at an increasing rate following acute exposure. Higher concentrations of nicotine led to a more pronounced effect. Even exposure to the pure basal fluid without nicotine resulted in a higher pulse rate. Having a chronically increased pulse rate is dangerous (Ganz, 1995; Umana, 2003). Tachycardia can result in a number of complications, including hypoxia, angina, syncope, and more (Mayo Clinic, 2022).

It has been established in the past that smoking traditional nicotine products also results in a substantial increase in resting heart rate via observational (Papathanasiou, 2013) and biological studies (Csordas, 2013). With this knowledge, it can be inferred that exposure to next-generation nicotine products would result in a similar rise in resting heart rate as well. However, a point of interest is the increase in heart rate with exposure to the basal e-liquid without nicotine, which can be observed in Figure 3. This shows that the basal fluid contained in e-cigarettes has cardiotoxic effects even without nicotine.

**Fig. 3.**
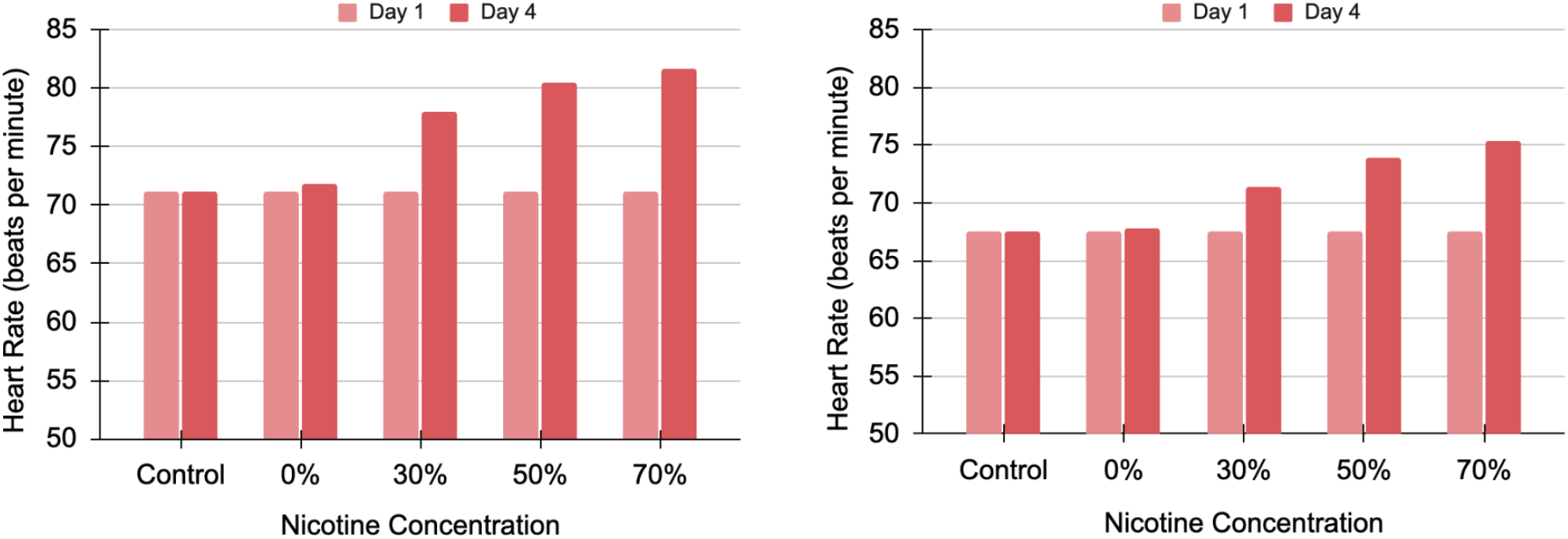
Heart rate trend over time for healthy cardiomyocytes (left) and diseased cells (right) with long QT syndrome. The data includes the control cells, cells treated with e-liquid basal fluid, and cells treated with 30%, 50%, and 70% nicotine concentration.

We found that acute exposure to e-liquid also caused cardiomyocytes to beat at a higher pulse pressure, according to the measurements conducted using the Sony SI8000 Cell Motion Imaging System. Pulse pressure can also be calculated by the difference between systolic and diastolic blood pressure. Thus, a higher pulse pressure is recognized as a risk factor for cardiovascular disease (Dart, 2001; de Simone, 2005). Generally, a pulse pressure exceeding 40 mmHg is considered unhealthy (Mayo Clinic 2022).

As shown in Figure 4, pulse pressure for the healthy cells after being treated with e-liquid exceeded well over 40 mmHg, indicating that smoking e-cigarettes may lead to hypertension. Once again, this finding is consistent with the already established connection between smoking cigarettes and malignant hypertension (Primatesta, 2001; Virdis, 2010). Complications of uncontrolled hypertension include aneurysm, myocardial infarction, stroke, and more (Sanyal, 2008; Mayo Clinic, 2022).

**Fig. 4.**
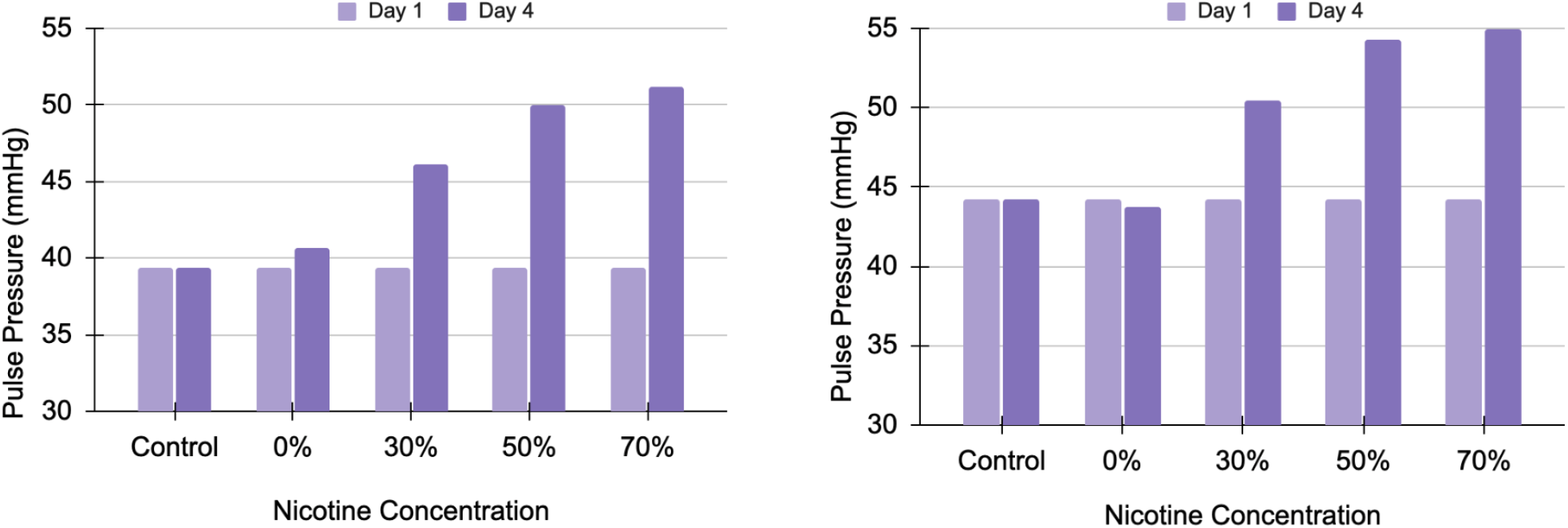
Pulse pressure trend over time for healthy cardiomyocytes (left) and diseased cells with long QT syndrome (right). The data includes the control cells, cells treated with e-liquid basal fluid, and cells treated with 30%, 50%, and 70% nicotine concentration.

The comparison of pulse pressure over time between the healthy cardiomyocytes and cardiomyocytes with long QT syndrome can be seen in Figure 4. Diseased cardiomyocytes were affected more drastically than healthy ones. The diseased cardiomyocytes with long QT syndrome have a higher starting pulse pressure (about 44 mmHg) relative to the healthy cardiomyocytes (about 39 mmHg). For example, at 70% nicotine concentration, healthy cells’ pulse pressure increased by to 51 mmHg whereas diseased cells’ pulse pressure increased to 59 mmHg. Overall, at the end of the 4-day testing period, with 30%, 50%, and 70% nicotine e-liquid exposure, the pulse pressure increased to 46 mmHg, 50 mmHg, and 51 mmHg for the healthy cardiomyocytes, and 51 mmHg, 54 mmHg, and 55 mmHg for the diseased cardiomyocytes, respectively. With this equivalent e-liquid exposure, both the healthy and the diseased cardiomyocytes experienced a similar magnitude of pulse pressure rise. However, the baseline pressure for the diseased cells was higher to begin with, which may result in life-threatening levels of pulse pressure. This result also correlates well with the fact that smoking is associated with higher mortality rates in people with congenital cardiovascular conditions (Engelfriet 2007; Gianicolo, 2010).

### B. Impact on Protein Concentration

Essential proteins, also known as the housekeeping proteins, are critical molecules for maintaining overall cell health. Through this work, we also found a direct correlation between acute exposure to e-liquid and degradation of essential proteins. The comparison between the healthy cardiomyocytes and cardiomyocytes with long QT syndrome can be seen in Figure 5. The protein concentration in the cardiomyocytes started at 77 ng/mL and 73 ng/mL for the healthy and diseased cells, respectively. These are in line with the 75-80 ng/mL range that is considered normal (ThermoFisher, 2021). The protein levels dropped to as low as 70 ng/mL in healthy cells and 66 ng/mL in diseased cells at the end of the trial. This aligns with the fact that cigarette smoke promotes ubiquitination and significant destabilization of the proteins (Kim, 2011; Han, 2017). Once again, higher levels of nicotine resulted in a more drastic degradation and the basal fluid was also detrimental to protein levels.

**Fig. 5.**
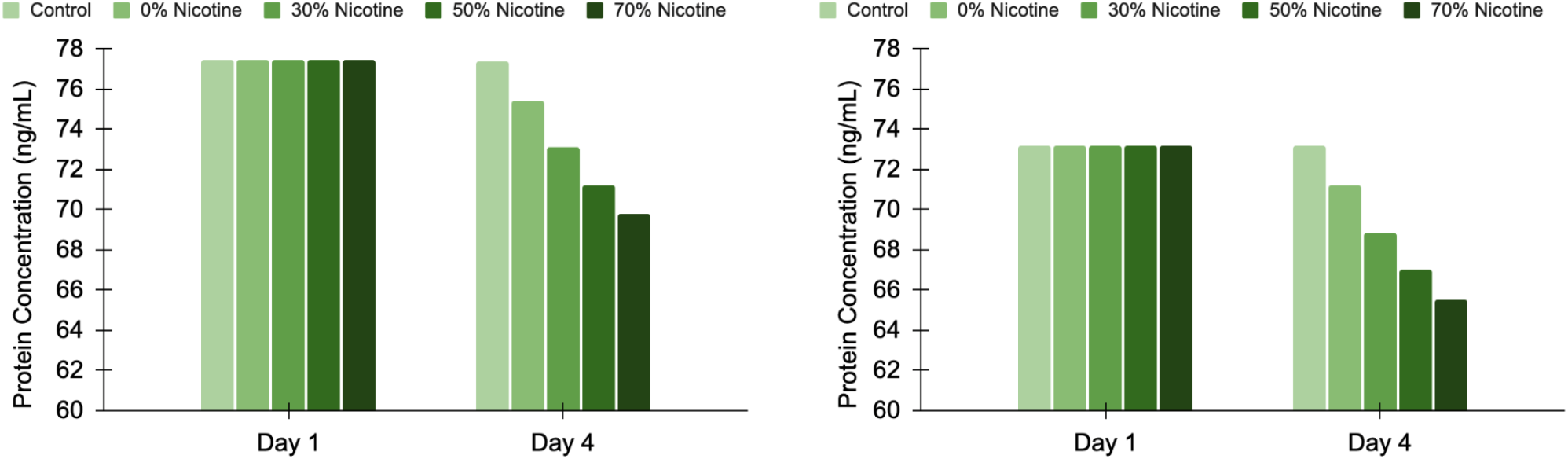
Protein concentration trend over time for healthy cardiomyocytes (left) and diseased cells with long QT syndrome (right). The data includes the control cells, cells treated with e-liquid basal fluid, and cells treated with 30%, 50%, and 70% nicotine concentration.

### C. Impact on Cardiomyocyte Viability

For the viability assay, we tracked the percentage of live cells over time. We utilized the ThermoFisher Cell Countess Chamber to check the amount of live vs. dead cells over the course of the experiment. We observed that the cell viability for both healthy and diseased cells declined over time following exposure to the e-liquid. The cardiomyocytes not only degraded in quality, but also began to die in large amounts. The comparison of viability between the healthy cardiomyocytes and cardiomyocytes with long QT syndrome can be seen in Figure 6. The higher viability recorded for 70% nicotine concentration in the right figure can be attributed to measurement variability. The control sample stayed at 100% viability, meaning that only a negligible amount of cells died naturally over the course of the testing period. Following e-liquid treatment, the cells began to die at increasingly rapid rates.

**Fig. 6.**
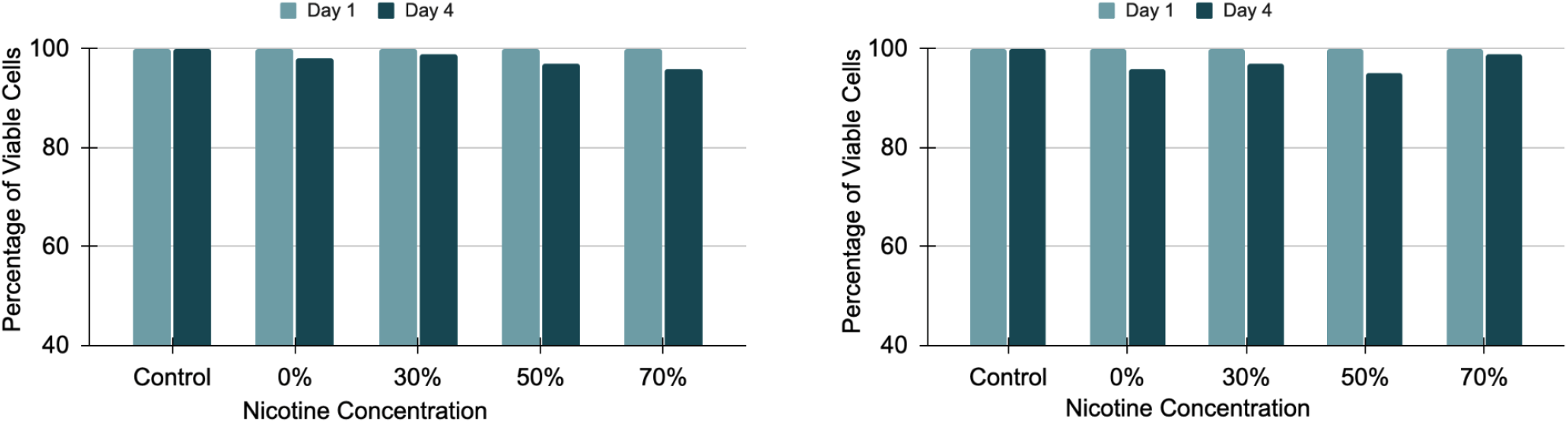
Viability trend over time for healthy cardiomyocytes (left) and diseased cells with long QT syndrome (right). The data includes the control cells, cells treated with e-liquid basal fluid, and cells treated with 30%, 50%, and 70% nicotine concentration.

## V. CONCLUSIONS

In summary, we have conducted an experimental study using cardiomyocytes derived from human induced pluripotent stem cells to analyze the effects of smoking e-cigarettes on the healthy and diseased human heart. We differentiated one healthy line of cells and one diseased line of cells with long QT syndrome into beating cardiomyocytes. We treated both cardiomyocyte lines with basal fluid and e-liquid with 30%, 50%, and 70% nicotine concentrations to determine the cardiotoxicity dependencies. After the acute exposure, we monitored and measured the condition of the cardiomyocytes over four days through three different functional assays.

Our studies demonstrated healthy and stable performances on key indicators from the controlled cell lines while acute exposure to e-liquid caused the conditions of the cardiomyocytes to deteriorate. Specifically, we observed an increase in the resting heart rate, a heightened pulse pressure, a decrease in the concentration of essential housekeeping proteins, as well as increased cell mortality. These are all risk factors for myocardial infarction, which can be lethal. The degradation of cardiomyocyte health is present when treated with pure basal fluid of the e-cigarettes and increases with higher nicotine concentrations.

While nicotine is the only cardiotoxic compound in regular cigarettes, e-cigarettes incorporate not only higher levels of nicotine but also a basal fluid. The increased nicotine concentration proportionally worsens the cardiomyopathic effects. Since e-cigarettes contain more nicotine in a single cartridge than 20 traditional cigarettes and there is no way for the users to track their own consumption, it is easier for the user to over-consume nicotine than in traditional tobacco products. The fact that the basal fluid exposure alone caused significant negative impacts to the cardiomyocytes indicates that the impact of e-cigarette is even worse than that of the traditional tobacco products.

Our work demonstrates the cardiotoxic effects of smoking e-cigarettes. Some potential ways to expand this study would be to pinpoint the exact mechanism in which e-liquid impacts cardiomyocytes by performing additional functional assays, such as quantitative polymerase chain reaction (q-PCR). Furthermore, it would be useful to determine the full scope of impact of e-cigarettes on the human body by carrying out similar experiments on parts of the brain, digestive tract, musculoskeletal system, and more.

## Notes

### Competing Interest Statement

The authors have declared no competing interest.

### Summary of Updates

Updated figures and references.

## REFERENCES

Bold KW, Kong G, Camenga DR, Simon P, Cavallo DA, Morean ME, Krishnan-Sarin S. (2018). Trajectories of e-cigarette and conventional cigarette use among youth. Pediatrics, 141 (1); e20171832. DOI: https://doi.org/10.1542/peds.2017-1832

Brown CJ, Cheng JM. (2014). Electronic cigarettes: product characterisation and design considerations. Tob Control, 23, ii4–ii10. DOI: 10.1136/tobaccocontrol-2013-051476

Burns DM, Lee L, Shen LZ, Gilpin E, Tolley HD, Vaugn J, Shanks TG. (1996). Cigarette smoking behavior in the United States. U.S. Department of Agriculture, Pages 13–42. https://cancercontrol.cancer.gov/sites/default/files/2020-08/m08_2.pdf

Burridge PW, Li YF, Matsa E, Wu H, Ong S, Sharma A, Holström A, Chang AC, Coronado MJ, Ebert AD, Knowles JW, Telli ML, Witteles RM, Blau HM, Bernstein D, Altman RB, Wu JC. (2016). Human induced pluripotent stem cell-derived cardiomyocytes recapitulate the predilection of breast cancer patients to doxorubicin-induced cardiotoxicity. Nat Med., 22(5), 547–556. DOI: 10.1038/nm.4087

Chatterjee K, Zhang J, Honbo N, Karliner JS. (2010). Doxorubicin cardiomyopathy. Cardiology, 115, 155–162. DOI: 10.1159/000265166

Cherian SV, Kumar A, Estrada-Y-Martin RM. (2020). E-cigarette or vaping product-associated lung injury: A review. The American Journal of Medicine, 133(6), 657–663. DOI: https://doi.org/10.1016/j.amjmed.2020.02.004

Collins, L. Glasser AM, Abudayyeh H, Pearson JL, Villanti AC. (2018). E-cigarette marketing and communication: How e-cigarette companies market e-cigarettes and the public engages with e-cigarette information. Nicotine &Tobacco Research, 21(1), 14–24. DOI: https://doi.org/10.1093/ntr/ntx284

Cherian SV, Kumar A, Estrada-Y-Martin RM. (2020). E-cigarette or vaping product-associated lung injury: A review. The American Journal of Medicine. 133(6), 657–663. DOI: https://doi.org/10.1016/j.amjmed.2020.02.004

Csordas A, Bernhard D. (2013). The biology behind the atherothrombotic effects of cigarette smoke. Nat Rev Cardiol., 10, 219–230. DOI: https://doi.org/10.1038/nrcardio.2013.8

Dart AM, Kingwell BA. (2001). Pulse pressure-a review of mechanisms and clinical relevance. J Am Coll Cardiol., 37(4) 975–984. DOI: doi.org/10.1016/S0735-1097(01)01108-1

de Simone G, Roman MJ, Alderman MH, Galderisi M, de Divitiis Oreste, Devereux RB. (2005). Is high pulse pressure a marker of preclinical cardiovascular disease? Am Journ Heart Assoc., 45, 575–579. DOI: https://doi.org/10.1161/01.HYP.0000158268.95012.08

Du P, Fan T, Yingst J, Veldheer S, Hrabovsky S, Chen C, Foulds J. (2019). Changes in e-cigarette use behaviors and dependence in long-term e-cigarette users. American Journal of Preventive Medicine, 57(3), 374–383. DOI: https://doi.org/10.1016/j.amepre.2019.04.021

Engelfriet PM, Drenthen W, Pieper PG, Tijssen JGP, Yap SC, Boersma E, Mulder BJM. (2008). Smoking and its effects on mortality in adults with congenital heart disease. International Journal of Cardiology, 127(1), 93–97, ISSN 0167-5273, DOI: https://doi.org/10.1016/j.ijcard.2007.05.008

Farsalinos KE, Polosa R, Cibella F, Niaura R. (2019). Is e-cigarette use associated with CHD and MI? Insights from the 2016 and 2017 National Health Interview Surveys. SAGE Journals. DOI: https://doi.org/10.1177/2040622319877741

Fermini B, Coyne ST, Coyne KP. (2018). Clinical trials in a dish: A perspective on the coming revolution in drug development. SLAS Discovery, 23(8), 765–776. DOI: https://doi.org/10.1177/2472555218775028

Ganz, LI, Friedman PL. (1995). Supraventricular tachycardia. N Engl J Med, 332, 162–173. DOI: 10.1056/NEJM199501193320307

Gianicolo AL, E., Monica C, Lamia AA, Ilenia F, Andreassi G M. (2010). Smoking and congenital heart disease: The epidemiological and biological link. Current Pharmaceutical Design, 16(23), 2572–2577.

Goldenberg I, Jonas M, Tenenbaum A. (2003). Current smoking, smoking cessation, and the risk of sudden cardiac death in patients with coronary artery disease. Arch Intern Med. 2301–2305. DOI: 10.1001/archinte.163.19.2301

Grskovic M, Javaherian A, Strulovici B, Daley GQ. (2013). Induced pluripotent stem cells– opportunities for disease modeling and drug discovery. Nature Reviews Drug Discovery, 10 (12), 915–29. DOI: 10.1038/nrd3577

Han S, Jerome JA, Gregory AD, Mallampalli RK. (2017). Cigarette smoke destabilizes NLRP3 protein by promoting its ubiquitination. Respir Res., 18, 2. DOI: https://doi.org/10.1186/s12931-016-0485-6

Hecht SS, Carmella SG, Kotandeniya D, Pillsbury ME, Chen M, Ransom BWS, Vogel RI, Thompson E, Murphy SE, Hatsukami DK. (2015). Evaluation of toxicant and carcinogen metabolites in the urine of e-cigarette users versus cigarette smokers. Nicotine &Tobacco Research, 17(6), 704–709, DOI: https://doi.org/10.1093/ntr/ntu218

Hudson L. (2022). Vaping statistics 2022. SingleCare, accessed December 24, 2022. https://www.singlecare.com/blog/news/vaping-statistics/

Karakikes I, Ameen M, Termglinchan V, Wu JC. (2015). Human induced pluripotent stem cell-derived cardiomyocytes. Circulation Research, 117(1), 80–88. DOI: https://doi.org/10.1161/CIRCRESAHA.117.305365

Kim SY, Lee JH, Huh JW, Ro JY, Oh YM, Lee SD, An S, Lee YS. (2011). Cigarette smoke induces Akt protein degradation by the ubiquitin-proteasome system. Journ of Bio Chem, 286, 37. DOI: https://doi.org/10.1074/jbc.M111.267633

Kitani T, Ong S, Lam C, Rhee J, Zhang JZ, Oikonomopoulos A, Ma N, Tian L, Lee J, Telli M, Witteles R, Sharma A, Sayed N, Wu JC. (2019). Human induced pluripotent stem cell model of Trastuzumab-induced cardiac dysfunction in breast cancer patients. Circulation, 139(21), 2451–2465. DOI: 10.1161/CIRCULATIONAHA.118.037357

Lee HW, Park SH, Weng MW, Wang HT, Huang WC, Lepor H, Wu XR, Chen LC, Tang MS. (2018). E-cigarette smoke damages DNA and reduces repair activity in mouse lung, heart, and bladder as well as in human lung and bladder cells. PNAS, 115(7), E1560–E1569. DOI: https://doi.org/10.1073/pnas.1718185115

Lee J, Termglinchan V, Diecke S, Itzhaki I, Lam CK, Garg P, Lau E, Greenhaw M, Seeger T, Wu H, Zhang JZ, Chen X, Gil IP, Ameen M, Sallam K, Rhee J, Churko JM, Chaudhary R, Chour T, Wang PJ, Snyder MP, Chang HY, Karakikes I, Wu JC. (2019). Activation of PDGF pathway links LMNA mutation to dilated cardiomyopathy. Nature, 572, 335–340. DOI: https://doi.org/10.1038/s41586-019-1406-x

Magdy T, Schuldt AJT, Wu JC, Bernstein D, Burridge PW. (2018). Human induced pluripotent stem cell (hiPSC)-derived cells to assess drug cardiotoxicity: Opportunities and problems. Annu Rev Pharmacol Toxicol, 58, 83–103. DOI: 10.1146/annurev-pharmtox-010617-053110

Matsa E, Rajamohan D, Dick E, Young L, Mellor I, Staniforth A, Denning C. (2011). Drug evaluation in cardiomyocytes derived from human induced pluripotent stem cells carrying a long QT syndrome type 2 mutation. European Heart Journal, 32(8), 952–962. DOI: https://doi.org/10.1093/eurheartj/ehr073

Matsa E, Burridge PW, Yu KH, Ahrens JH, Tennglinchan V, Wu H, Liu C, Shukla P, Sayed N, Churko JM, Shao N, Woo NA, Chao AS, Gold JD, Karakikes I, Snyder MP, Wu JC. (2016). Transcriptome profiling of patient-specific human iPSC-Cardiomyocytes products individual drug safety and efficacy responses in vitro. Cell Stem Cell, 19(3), 311–325. DOI: 10.1016/j.stem.2016.07.006

Mattingly DT, Hirschtick JL, Meza R, Fleischer NL. (2020). Trends in prevalence and sociodemographic and geographic patterns of current menthol cigarette use among U.S. adults. Preventive Medicine Reports, 20, 101227. DOI: https://doi.org/10.1016/j.pmedr.2020.101227

Mayo Clinic. (2022). Tachycardia. Accessed December 25, 2022. https://www.mayoclinic.org/diseases-conditions/tachycardia/symptoms-causes/syc-20355127

Mayo Clinic. (2022). Pulse pressure: An indicator of heart health? Accessed December 25, 2022. https://www.mayoclinic.org/diseases-conditions/high-blood-pressure/expert-answers/pulse-pressure/faq-20058189#.

Mayo Clinic. (2022). High blood pressure (hypertension). Accessed December 25, 2022. https://www.mayoclinic.org/diseases-conditions/high-blood-pressure/symptoms-causes/syc-20373410

Narkar A, Willard JM, Blinova K. (2022). Chronic cardiotoxicity assays using human induced pluripotent stem cell-derived cardiomyocytes (hiPSC-CMs). Int J Mol Sci, 23(6), 3199. doi: 10.3390/ijms23063199.

Ottervanger JP, Festen JM, de Vries AG, Stricker BHC. (1995). Acute myocardial infarction while using the nicotine patch. CHEST, 107(6), 1765–1766. DOI: https://doi.org/10.1378/chest.107.6.1765

Papathanasiou G, Georgakopoulos D, Papageorgiou E, Zerva E, Michalis L, Kalfakakou V, Evangelou A. (2013). Effects of smoking on heart rate at rest and during exercise, and on heart rate recovery, in young adults. Hellenic J Cardiol, 54(3), 168–77. https://pubmed.ncbi.nlm.nih.gov/23685653/

Pillai-Kastoori L, Schutz-Geschwender AR, Harford JA. (2020). A systematic approach to quantitative Western blot analysis. Anal Biochem, 593, 113608. DOI: 10.1016/j.ab.2020.113608

Primatesta P, Falaschetti E, Gupta S, Marmot MG, Poulter NR. (2001). Association between smoking and blood pressure. J Am Heart Assoc, 37, 187–193. DOI: https://doi.org/10.1161/01.HYP.37.2.187

Qasim H, Karim Z, Rivera J, Khasawneh F, Aishbool F. (2017). Impact of electronic cigarettes on the cardiovascular system. J Am Heart Assoc, 6, e006353. DOI: https://doi.org/10.1161/JAHA.117.006353

Ramamurthi D, Chau C, Jackler R. (2021). Exploitation of the COVID-19 pandemic by ecigarette marketers. Tobacco Control, 30, e56–e59. DOI: http://dx.doi.org/10.1136/tobaccocontrol-2020-055855

Rowell TR, Tarran R. (2015). Will chronic e-cigarette use cause lung disease? American Journal of Physiology, 309(12), L1398–L1409. DOI: https://doi.org/10.1152/ajplung.00272.2015

Sanyal AJ, Bosch J, Blei A, Arroyo V. (2008). Portal hypertension and its complications. Gastroenterology, 134, 1715–1728. DOI: https://doi.org/10.1053/j.gastro.2008.03.007

Shi Y, Inoue H, Wu JC, Yamanaka S. (2017). Induced pluripotent stem cell technology: a decade of progress. Nat Rev Drug Discov, 16, 115–130 https://doi.org/10.1038/nrd.2016.245

Smith CJ, Osborn AM. (2009). Advantages and limitations of quantitative PCR (q-PCR)-based approaches in microbial ecology. FEMS Microbiology Ecology, 67(1), 6–20. DOI: https://doi.org/10.1111/j.1574-6941.2008.00629.x

Sun T, Lim CCW, Stjepanovič D, Leung J, Connor JP, Gartner C, Hall WD, Chan GCK. (2021). Has increased youth e-cigarette use in the USA, between 2014 and 2020, changed conventional smoking behaviors, future intentions to smoke and perceived smoking harms? Addictive Behaviors. DOI: https://doi.org/10.1016/j.addbeh.2021.107073

Takahashi K, Yamanaka S. (2006). Induction of pluripotent stem cells from mouse embryonic and adult fibroblast cultures by defined factors. Cell Press, 126(4), 663–676. DOI: https://doi.org/10.1016/j.cell.2006.07.024

ThermoFisher. (2021). Quantitative western blot analysis. Accessed December 31, 2022. https://www.thermofisher.com/us/en/home/life-science/protein-biology/protein-biology-learning-center/protein-biology-resource-library/pierce-protein-methods/quantitative-western-blot-analysis.html#>.

Umana E, Solares CA, Alpert MA. (2003). Tachycardia-induced cardiomyopathy. Am Journ of Med, 114(1), 51–55. DOI: https://doi.org/10.1016/S0002-9343(02)01472-9

Virdis A, Giannarelli C, Fritsch MN, Taddei S, Ghiadoni L. (2010). Cigarette smoking and hypertension. Current Pharmaceutical Design, 16(23), 2517–2525.

Wei T, Chandy M, Nishiga M, Zhang A, Kumar KK, Thomas D, Manhas A, Rhee S, Justesen JM, Chen IY, Wo H, Khanamiri S, Yang JY, Seidi FJ, Burns NZ, Liu C, Sayed N, Shie J, Yeh C, Yang K, Lau E, Lynch KL, Rivas M, Kobilka B, Wu JC. (2022). Cannabinoid receptor 1 antagonist genistein attenuates marijuana-induced vascular inflammation. Cell, 185(10), 1676–1693.e23. DOI: 10.1016/j.cell.2022.04.005

Ye L, Swingen C, Zhang J. (2013). Induced pluripotent stem cells and their potential for basic and clinical sciences. Current Cardiology Reviews, 9(1), 63–72. DOI: 10.2174/157340313805076278

